# The *Pseudomonas aeruginosa* PilSR two-component system regulates both twitching and swimming motilities

**DOI:** 10.1101/323923

**Authors:** Sara L.N. Kilmury, Lori L. Burrows

## Abstract

Motility is an important virulence trait for many bacterial pathogens, allowing them to position themselves in appropriate locations at appropriate times. Motility structures - pili and flagella - are also involved in sensing surface contact, which modulates pathogenicity. In *Pseudomonas aeruginosa*, the PilS-PilR two-component system (TCS) regulates expression of the type IV pilus (T4P) major subunit PilA, while biosynthesis of the single polar flagellum is regulated by a hierarchical system that includes the FleSR TCS. Previous studies in *Geobacter sulfurreducens* and *Dichelobacter nodosus* implicated PilR in regulation of non-T4P-related genes, including some involved in flagellar biosynthesis. Here we used RNAseq analysis to identify genes in addition to *pilA* with changes in expression in the absence of *pilR*. Among these were 10 genes inversely dysregulated by loss of *pilA* versus *pilR*, even though both *pilA* and *pilR* mutants lack T4P and pilus-related phenotypes. The products of those genes - many of which were hypothetical - may be important for virulence and surface-associated behaviours, as mutants had altered swarming motility, biofilm formation, type VI secretion, and pathogenicity in a nematode model. Further, the PilSR TCS positively regulated transcription of *fleSR*, and thus many genes in the FleSR regulon. As a result, *pilSR* deletion mutants had defects in swimming motility that were independent of the loss of PilA. Together these data suggest that in addition to controlling T4P expression, PilSR have a broader role in the regulation of *P. aeruginosa* motility and surface sensing behaviours.

## IMPORTANCE

Surface appendages like type IV pili and flagella are important for establishing surface attachment and infection in a host in response to appropriate cues. The PilSR regulatory system that controls type IV pilus expression in *Pseudomonas aeruginosa* has an established role in expression of the major pilin PilA. Here we provide evidence supporting a new role for PilSR in regulating flagellum-dependent swimming motility in addition to pilus-dependent twitching motility. Further, even though both *pilA* and *pilR* mutants lack PilA and pili, we identified sets of genes downregulated in the *pilR* mutant and upregulated in a *pilA* mutant as well as those downregulated only in a *pilR* mutant, independently of pilus expression. This finding suggests that change in the inner membrane levels of PilA is only one of the cues to which PilR responds to modulate gene expression. Identification of PilR as a regulator of multiple motility pathways may make it an interesting therapeutic target for anti-virulence compounds.

## INTRODUCTION

Prokaryotes rely on the use of two-component regulatory systems (TCS) to control many of their cellular activities. Typically comprised of a membrane-bound histidine sensor kinase and a cytoplasmic response regulator, TCSs allow bacteria to respond rapidly to chemical and physical changes in their intra-or extracellular environments, altering expression of specific genes in response to a stimulus (1). The opportunistic pathogen *Pseudomonas aeruginosa* encodes a higher-than-average number of TCSs (2) that control diverse functions, including several motility phenotypes. Flagellum-dependent swimming motility, for example, is controlled through a regulatory cascade that includes the transcriptional regulator, FleQ (3) and the FleS-FleR TCS, which like many TCSs also requires the alternate sigma factor RpoN (σ^54^)(4). FleQ controls transcription of *fleS-fleR* in addition to multiple other flagellar, adhesion and biofilm-associated genes, in a c-di-GMP dependent manner (3,5). FleSR has been implicated in the expression of 20 or more flagellar biosynthetic genes in *P. aeruginosa*, as well as additional genes not previously known to be involved in flagellar assembly or function (6).

The other major motility system in *P. aeruginosa* is the type IV pilus (T4P) system, which is used for twitching across solid and semi solid surfaces (7,8) among other important functions. In contrast to the single polar flagellum that is used to propel the cell in low viscosity media, the cell extends multiple pili that retract - independently or in a coordinated bundle - pulling it towards the point of attachment (9–11). Pili can be extended from either pole, but typically a single pole is used at one time allowing for directional movement (12). The majority of the pilus fibre is made of hundreds to thousands of subunits of the major pilin protein, PilA (13), the expression of which could be energetically costly to the cell if not tightly controlled.

*pilA* transcription is regulated by another TCS, PilS-PilR, in *P. aeruginosa* and many other T4P-expressing bacteria (14–18). PilS is an atypical sensor histidine kinase (SK) with 6 transmembrane segments (19–21) that allow PilS to interact directly with PilA for pilin autoregulation (22). PilR is the cytoplasmic response regulator (RR) that binds in conjunction with σ^54^ to the *pilA* promoter to activate transcription (23,24). Neither *pilA* nor *pilR* mutants express PilA and therefore T4P, but they have opposite PilR activation states. Activation of PilR upon transient decreases in PilA levels may be one way in which pilus attachment events are detected by the cell.

In contrast to the response regulator FleR, which has a well-defined regulon in *P. aeruginosa* (6), the suite of genes potentially controlled by PilR is poorly characterized. Genetic and *in silico* analyses of the PilR regulons of *Geobacter sulfurreducens* (16,25) and *Dichelobacter nodosus* (17) have been performed, but comparable studies are lacking in *P. aeruginosa*. Screening of the *G. sulfurreducens* genome for putative PilR binding sites revealed 54 loci with hypothesized σ^54^-dependent promoters, many of which were upstream of genes for T4P and flagellar biosynthesis, or cell wall biogenesis (25). Those data, in combination with work performed in *D. nodosus*, which identified several surface-exposed proteins whose expression was controlled by PilR (17), suggest that *P. aeruginosa* PilR likely has additional functions beyond control of *pilA* transcription. However, each of the cited studies focused mainly on identification of genes and characterization of their pilus-related functions without examining other phenotypic consequences of loss of *pilR*.

In this work, we used RNAseq analysis to identify genes that were dysregulated by loss of *pilR*. Because *pilR* mutants lack pili, which are important for surface sensing (26) and control of downstream events such as biofilm formation (27), we included a *pilA* mutant in our analysis to distinguish genes whose expression is specifically controlled by PilR from those that are affected by the loss of PilA. In addition to several genes that were co-regulated with *pilA*, which we have termed “pilin-responsive” genes, we also identified multiple flagellar genes, including the FleSR TCS, as being downregulated only in the absence of *pilR*, in a pilin unresponsive manner. We show that the consequence of this downregulation is a previously unreported defect in swimming motility in both *pilS* and *pilR* mutants, independent of the loss of PilA. This work defines the pilin-dependent and independent regulons of PilR and provides evidence for a direct regulatory connection between the *P. aeruginosa* T4P and flagellar motility systems.

## RESULTS

### The expression of multiple genes is similarly altered in pilA and pilR mutants

We performed RNAseq analysis to identify genes in addition to *pilA* that might be controlled by the PilSR TCS. However, in designing this experiment we considered that i) *pilR* mutants also lack expression of PilA; ii) loss of PilA contributes to a decrease in intracellular levels of the messenger molecule cyclic adenosine monophosphate (cAMP) (28); and iii) there are over 200 genes in *P. aeruginosa* that are at least partially cAMP-dependent, including Vfr, a cAMP-binding virulence factor regulator (28). To separate genes that are affected by the loss of PilA that occurs in both *pilA* and *pilR* mutants from those that are truly regulated by PilR, we categorized genes as those whose expression was changed in only the *pila* or *pilR* backgrounds, versus both backgrounds, compared to WT PAK (**Figure 1**). The former group may also include genes that are cAMP-dependent. We did not include a *pUS* mutant in RNAseq analysis, because PilS potentially interacts with alternate response regulators, making it more challenging to distinguish genes that are controlled by PilSR and those regulated by PilS and other unidentified RRs (29). Genes that were dysregulated similarly by at least 2-fold in both the *pilA* and *pilR* mutants are summarized in **Table S1**, and included several T4P-associated genes such as *tsaP* (30), and minor pilins *fimU, pilV, pilW, pilY1* and *pilE* (31), previously identified as being Vfr dependent (28). In total, 18 of 56 genes in this category (highlighted in gray in **Table S1)** are also Vfr and cAMP-dependent (28). Since the expression of genes in this class was affected by loss of PilA, suggesting PilR’s role is indirect, they were not examined further. No genes whose expression was dysregulated by loss of *pilA* but not *pilR* were identified.

**Figure 1.**
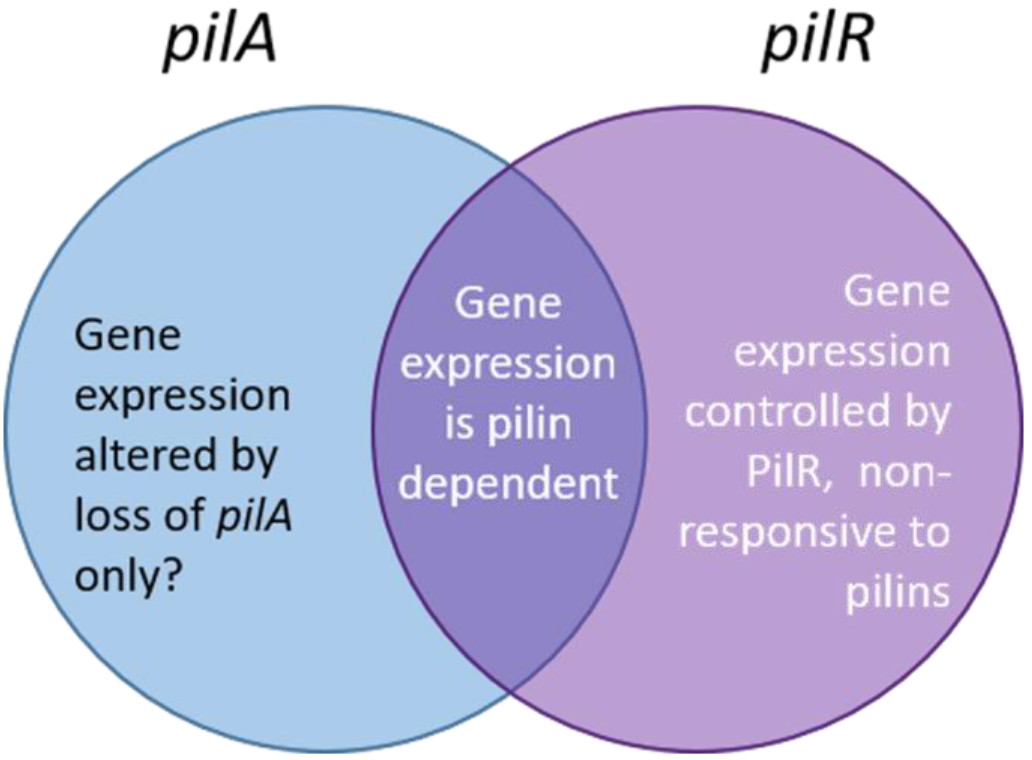
RNAseq experimental design. Our RNAseq experiment was designed to distinguish between genes dysregulated by loss of *pilR* versus loss of *pilA*, as *pilR* mutants also lack the pilin protein. Genes coordinately dysregulated in both *pilA* and *pilR* compared to WT may be due to loss of PilA or pilus expression. Genes that are inversely dysregulated in *pilA* and *pilR* mutants are upregulated in the absence of PilA (pilin responsive), in a *pilR-dependent* manner. Genes dysregulated by loss of *pilR* but not *pilA* are dependent only on PilR expression and referred to as pilin unresponsive in this study

### Ten genes are inversely dysregulated by loss of pilA versus pilR

The expression of a subset of ten genes was decreased in the *pilR* mutant but markedly increased in the *pilA* mutant, even though *pilR* mutants also lack PilA (23) (**Figure 2**, **Table S2**). We categorized these genes as ‘pilin-responsive’, because similar to PilA, their expression was dependent on PilR and increased when PilA levels were low. All but 3 of these genes encode hypothetical proteins or are unannotated in the PAO1 genome. The co-regulation of these genes with *pilA* suggests that their products could be previously unidentified contributors to T4P biogenesis and/or function, or to other forms of motility.

**Figure 2:**
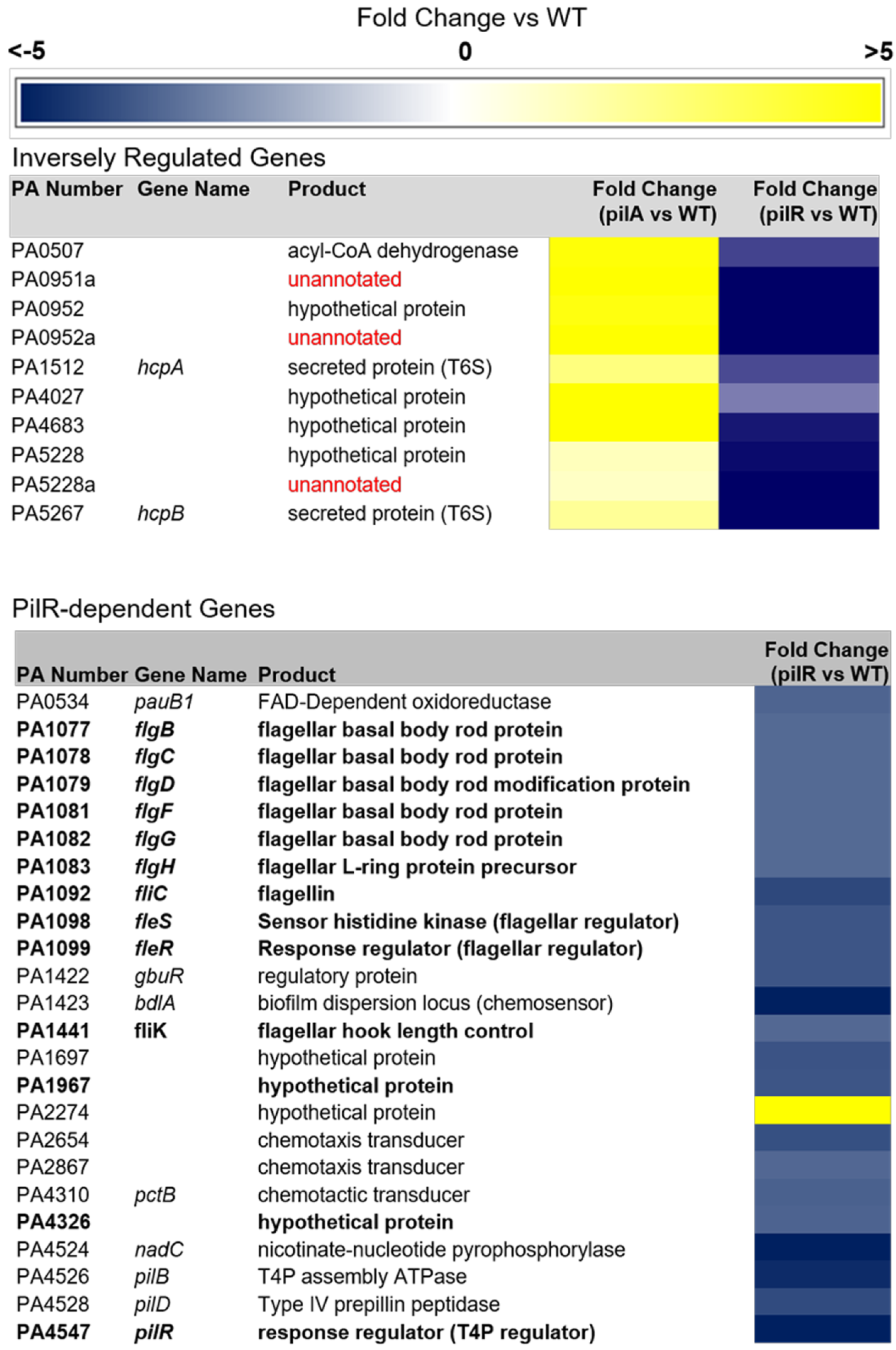
Multiple T4P and non-T4P genes are dysregulated by loss of *pilR*. The heat maps highlight select genes of interest that are >3-fold dysregulated by the loss of *pilR*, or both *pilA* and *pilR*. In addition to *pilA*, several other T4P associated genes have altered expression in *pilA*and/or *pilR* mutants. Expression of ten genes, mostly hypothetical or unannotated, is increased in a *pilA* mutant but decreased in *pilR*, which also lacks PilA. Of the genes dysregulated in a *pilR* mutant, many were associated with flagellar biosynthesis and function, including those encoding the FleS-FleR TCS.

To test this hypothesis, we extracted mutants with insertions in homologs of those PAK genes from the ordered PA14 transposon (Tn) library (32). There were no transposon insertions in three of the ten genes, and one additional mutant failed to grow in liquid culture. The PAO1 and PA14 designations for the remaining 6 genes of interest for which mutants were available are listed in **Table 1.** We tested these mutants for twitching, swimming, and swarming motilities. While all had wild-type twitching motility, insertions in PA14_51940 (PA0952), PA14_11740 (PA4027) and PA14_69560 (PA5267, *hcpB*) caused defects in swarming. Disruption of PA14_695060 (*hcpB*) also reduced swimming, alluding to a role in flagellar function or biosynthesis, in addition to its established function in Type VI secretion **(Figure 3).** Together, these data indicate that genes co-regulated with *pilA* are not necessarily required for T4P function, but a subset are involved in other forms of motility and in some cases, biofilm formation (33) and pathogenicity in *C. elegans* (34) (**Figure SI**).

**Figure 3:**
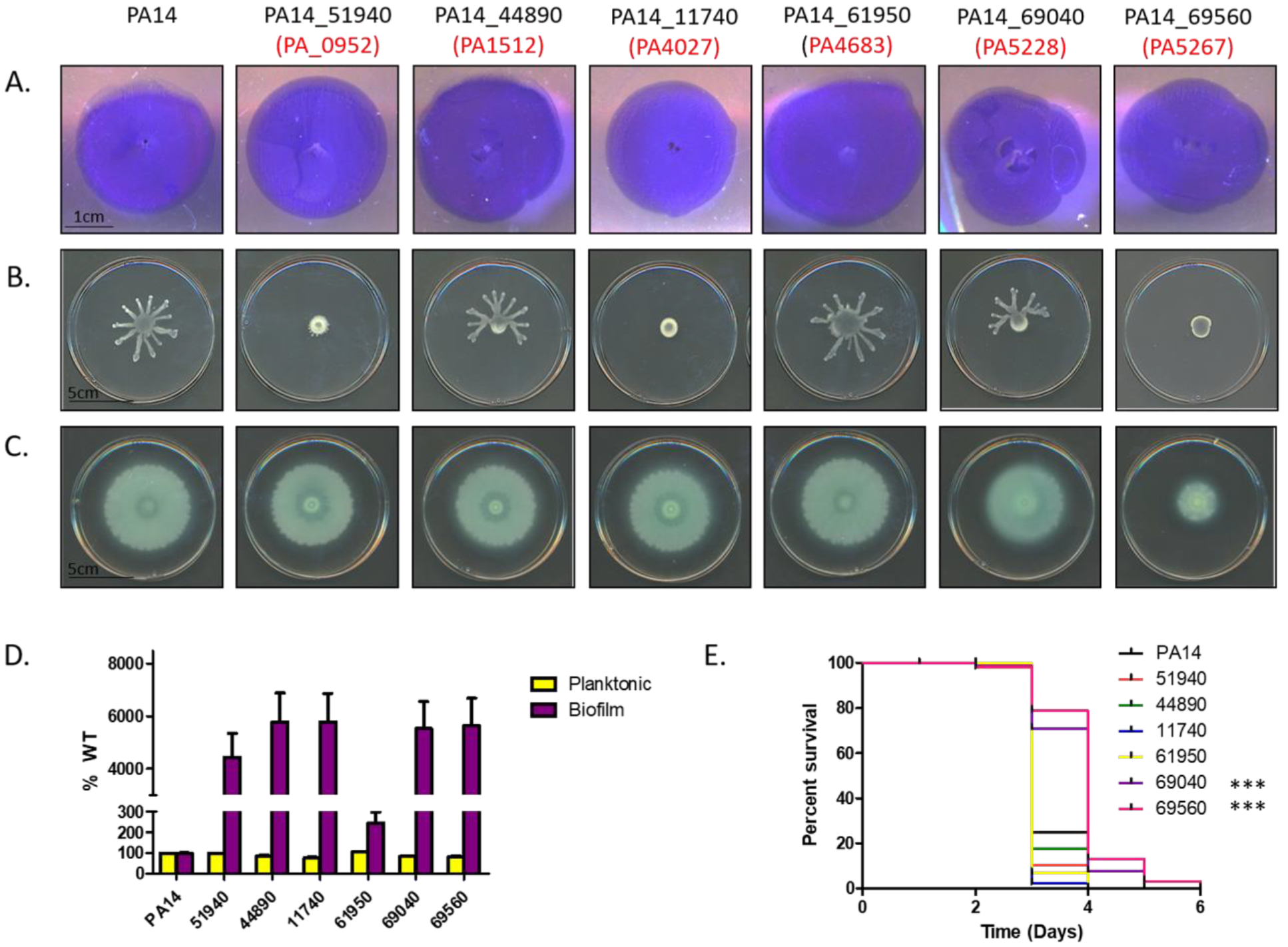
PA14 mutant homologs of inversely dysregulated genes affect motility phenotypes. Available PA14 transposon mutant homologs of inversely dysregulated genes identified in PAK were tested for **A)** twitching, **B)** swarming and **C)** swimming motilities, **D)** biofilm formation, and **E)** pathogenicity towards *Caenorhabditis elegans* to identify the functions of hypothetical proteins. All mutants displayed WT twitching, but some had varying defects in swimming and/or swarming motility. Several mutants also exhibited a hyperbiofilm phenotype, while two had defects in *C. elegans* killing (*** p<0.005).

**Table 1.**
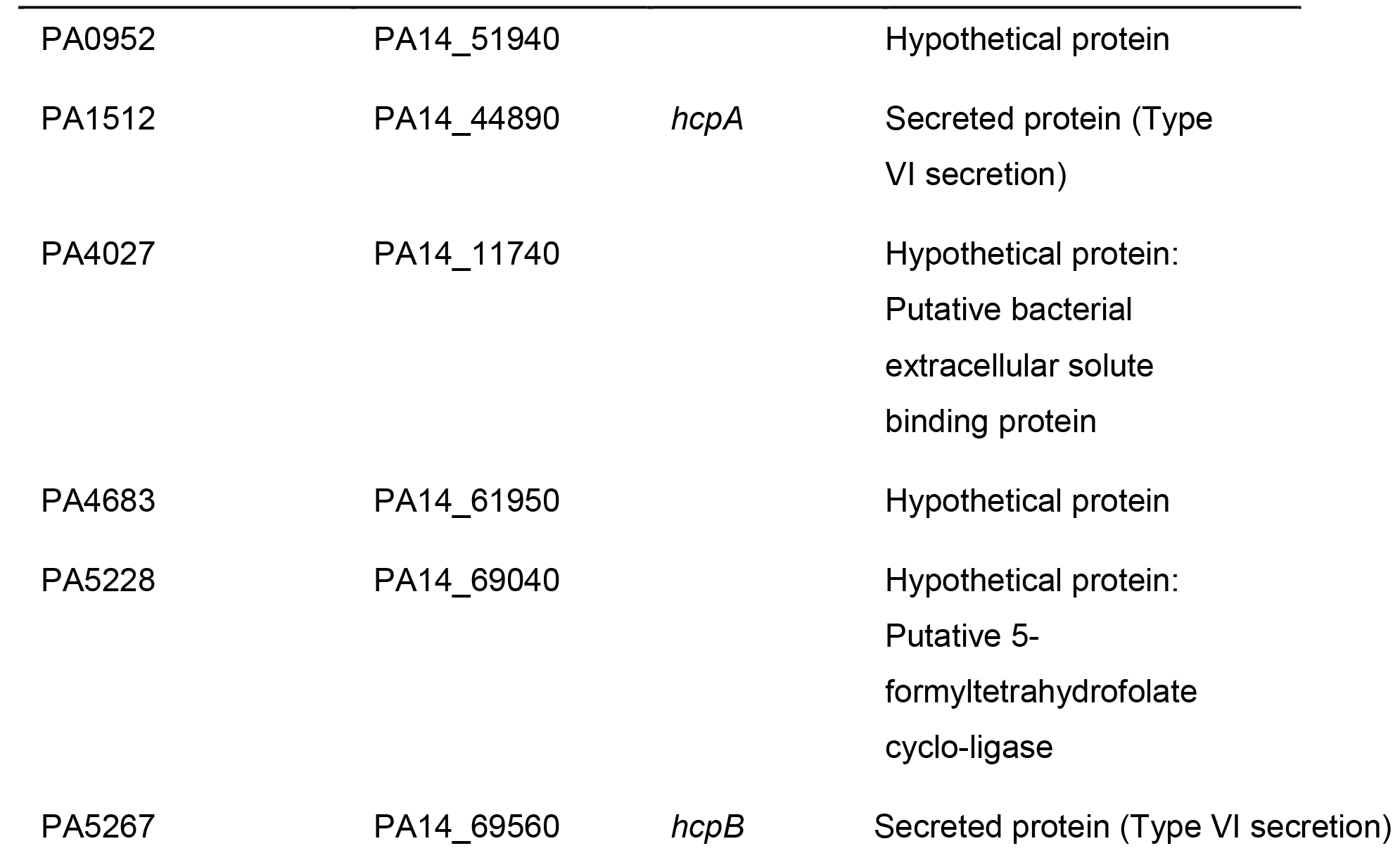
**PA and corresponding PA14 gene numbers of inversely dysregulated (pilin responsive) genes**

**Table 2.** **Table 2. Strains and plasmids used in this study**

| **Strain** | **Description** | **Source** |
| --- | --- | --- |
| PAK WT | WT Group II strain of *P. aeruginosa* | (J. Boyd) |
| *pilA* | PAK with chromosomal deletion of *pilA* | (This study) |
| *pilS* | PAK with chromosomal deletion of *pilS* | (This study) |
| *pilR* | PAK with chromosomal deletion of *pilR* | (This study) |
| *fliC* | PAK with FRT insertion in *fliC* | (This study) |
| *fleSR* | PAK with chromosomal of the full *fleS-fleR operon* | (This study) |
| PA14 WT | WT Group III strain of *P. aeruginosa* | (32) |
| PA14_51950 | PA14 with a transposon insertion in gene PA14_51950 | (32) |
| PA14_44890 | PA14 with a transposon insertion in PA14_44890 (*hcpA*) | (32) |
| PA14_11740 | PA14 with a transposon insertion in PA14_11740 | (32) |
| PA14_61950 | PA14 with a transposon insertion in PA14_61950 | (32) |
| PA14_69040 | PA14 with a transposon insertion in PA14_69040 | (32) |
| PA14_69560 | PA14 with a transposon insertion in PA14_69560 (*hcpB*) | (32) |
| **Plasmid** |  |  |
| pMS402-*ppilA* | *pilA* promoter cloned into the BamHI site of pMS402, putting lux genes under control of *pilA* promoter | (22) |

### A subset of genes is dysregulated only by loss of PilR

In the *pilR* mutant, 89 genes were dysregulated ≥2-fold **(Table S3).** To prioritize our follow-up studies, we focused on a shorter list of genes with ≥3-fold changes in expression. Prior to this study, *pilA* was the only known member of the PilS-PilR regulon in *P. aeruginosa* (19), though studies in *G. sulfurreducens* and *D. nodosus* suggested its regulon was likely to be broader (16,17, 25). Of particular interest were 24 genes whose expression was ≥3-fold altered in *pilR* mutants but unaffected by loss of *pilA*. These *pilR-*dependent but pilin unresponsive genes are highlighted in **Figure 2.** According to the *Pseudomonas* genome database (35), these genes include five putative chemotactic transducers, two biofilm-associated chemosensory proteins, six hypothetical proteins, and several metabolic enzymes. However, motility-associated genes were the most common class identified. The genes encoding the T4P assembly ATPase, PilB and prepilin peptidase, PilD, which share a divergently oriented promoter with *pilA*, were downregulated in *pilR* but unaffected by loss of *pilA* (**Figure 2**), even though previous studies suggested they were controlled by σ^70^, not PilSR and σ^54^ (36).

### Multiple flagellum biosynthetic genes are downregulated in a pilR mutant

In addition to the T4P-associated genes above, several flagellum biosynthetic genes had decreased expression only in the *pilR* background **(Figure 2, Table S3 bolded text).** Among them were *fleS-fleR* encoding the FleSR TCS, part of a regulatory cascade that controls the expression of genes associated with flagellum biosynthesis and function (4,6). Each had approximately 3-fold lower expression in *pilR* compared to WT, while there was no difference in their expression in *pilA* versus WT. This trend was verified using RT-PCR, though the magnitude was closer to 2-fold by this method (Supplemental **Figure S2).** Of the flagellar genes in this category (**Figure 2**), 10 of 12 (excluding *fleS* and *fleR*) are *fleR* dependent (6). The remaining two, *fliE* and *fliF*, are FleQ dependent, but also had decreased (≥2-fold) transcription in *a fleR* mutant in a previous study (6). These data suggest that PilSR positively regulates *fleSR* expression, and when PilR is absent, expression of FleSR-dependent genes is decreased accordingly.

### Swimming motility is impaired by loss of pilS-pilR

We next tested if downregulation of *fleSR* in the *pilS* and *pilR* backgrounds impacted swimming motility, using a plate-based assay. A*fliC* mutant lacking the flagellin subunit was used as a negative control. *pilA* mutants swam comparably to WT PAK, while *pilS* and *pilR* mutants - which also lack surface pili - exhibited significant swimming defects (p<0.005), with uniform zones that reached about 40% of WT (**Figure 4, dashed line.**) Interestingly, both *pilS* and *pilR* mutants produced flares with increased motility extending beyond these uniform swimming zones. These flares were hypothesized to be the result of suppressor mutations that could overcome the effect of *pila* or *pilR* deletion on swimming.

**Figure 4:**
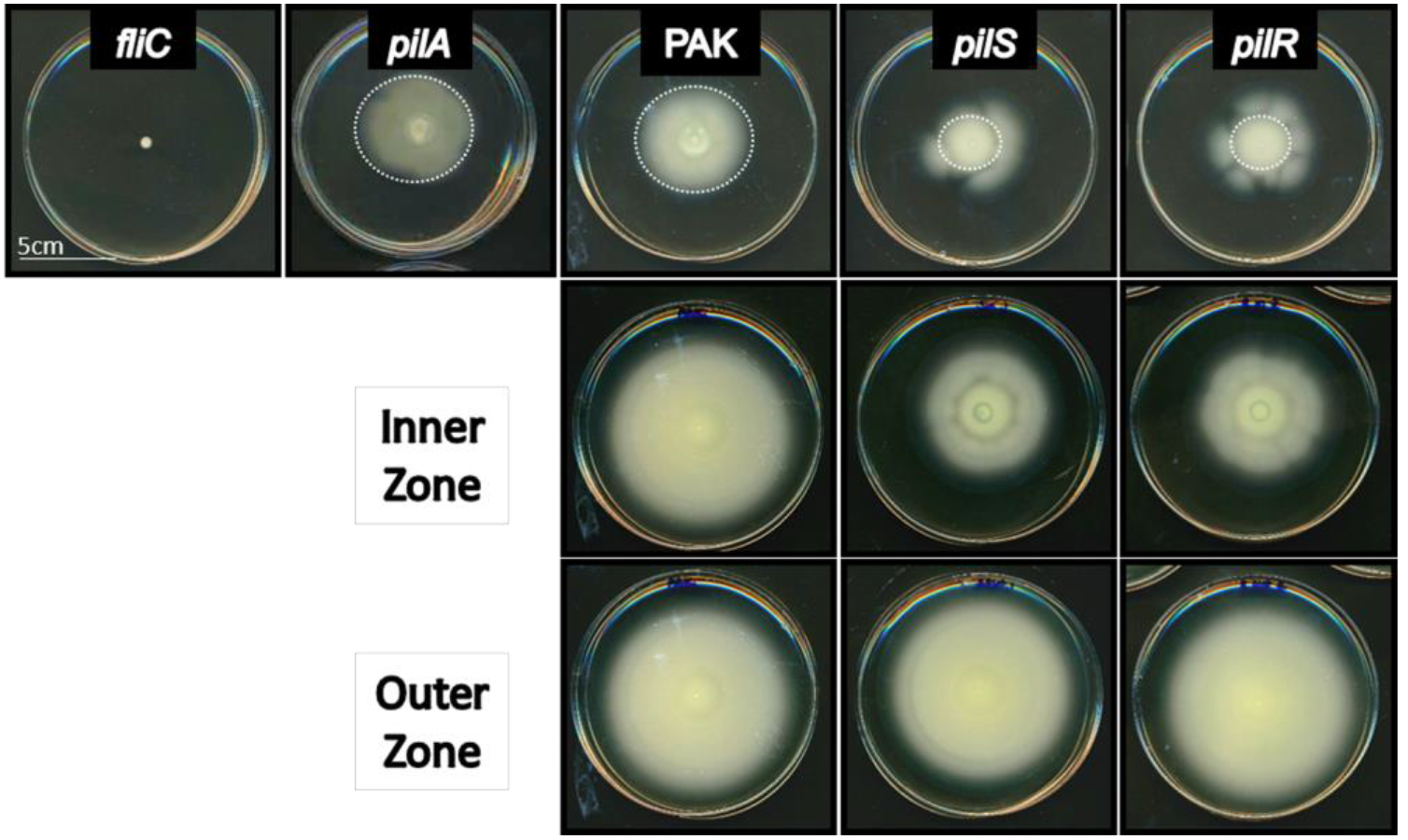
Swimming motility is impaired in *pilS* and *pilR* mutants. Loss of *pilS* or *pilR* results in decreased swimming motility (~40% of WT) in a plate-based assay. *pilA* mutants swim comparably to WT, indicating that the swimming defect is not PilA dependent. *pilS* and *pilR* mutants appear to acquire suppressors that overcome these defects resulting in asymmetrical flares. Reinoculation of swimming plates with cells from the interior of swimming zones—inside the dotted white circles of *pilSR* (inner) recapitulate the original phenotype, while cells taken from the flares (outer, from flares outside the white circle, except for WT) swim to WT levels. Asterisks denote the location from which cells were taken for the reinoculated swimming plates.

To test this idea, we isolated cells from the inner swimming zones of *pila* and *pilR* plates (inside the dashed line, **Figure 4**) and the putative suppressor mutants (flares outside the dashed line) and reassessed their ability to swim after culturing them overnight. As controls, we took samples from the WT zone close to the point of inoculation (‘inner’) and from the outer edge of the swimming zone (‘oute’). Repeating the swimming assays with these samples revealed no difference in swimming between inner and outer samples from WT. However, p/75 and *pilR* cells taken from the inner swimming zones recapitulated the original swimming motility defects of the mutants - including the re-appearance of highly motile suppressors - while cells taken from the outer flares had motility comparable to WT (**Figure 4**), indicating that they likely acquired mutation(s) that allow for full motility in the absence of *pilSR*.

To test if other flagellum-dependent phenotypes were affected by loss of *pilSR*, we measured swarming motility, using the original mutants and the suppressors isolated from the swimming experiments above. *pilSR* mutants in PAO1 were previously reported to be non-swarmers (37), but in our hands the same mutants in the PAK background retain partial swarming motility, albeit with an altered morphology compared to WT. The PAK *pilSR* mutants swarmed similarly to a *pilA* mutant, with fewer and irregular tendrils (**Figure S2**). Interestingly, *pilSR* mutants isolated from the outer flares of the swimming plates in **Figure 4** had swarming motility comparable to those isolated from the inner zones and the parent *pilS* and *pilR* strains. While flagella are required for swarming, the suppressor mutations that restored swimming motility in the *pilS* and *pilR* backgrounds did not restore swarming, suggesting that expression of distinct swarming-related genes remains dysregulated.

### FleSR impact twitching motility and pilA expression

RNAseq analyses revealed that PilR was required for wild type expression of *fleSR*. We next tested if this was a reciprocal regulatory pathway in which FleSR might contribute to regulation of *piIS*-*pilR* and the PilSR regulon. We tested if loss of *fleSR* affected *pilA* expression and/or T4P function. A double deletion *of fleSR* was made in the PAK background, and twitching motility measured. Loss of *fleSR* reduced twitching motility to a modest but significant extent (p<0.005), with the double mutant reproducibly twitching to approximately 80% of WT (**Figure 5A**). Interestingly, when *pilA* transcription was monitored using a *lux-pilA* reporter assay, *fleS-fleR* mutants had increased *pilA* transcription compared to WT over a 5 h time course (**Figure 5B**). Therefore, while FleS-FleR are involved in the modulation of twitching motility and *pilA* transcription, it is not yet clear if this occurs directly through regulation of *pilSR*, as increased levels of PilA can inhibit PilSR activation (22).

**Figure 5:**
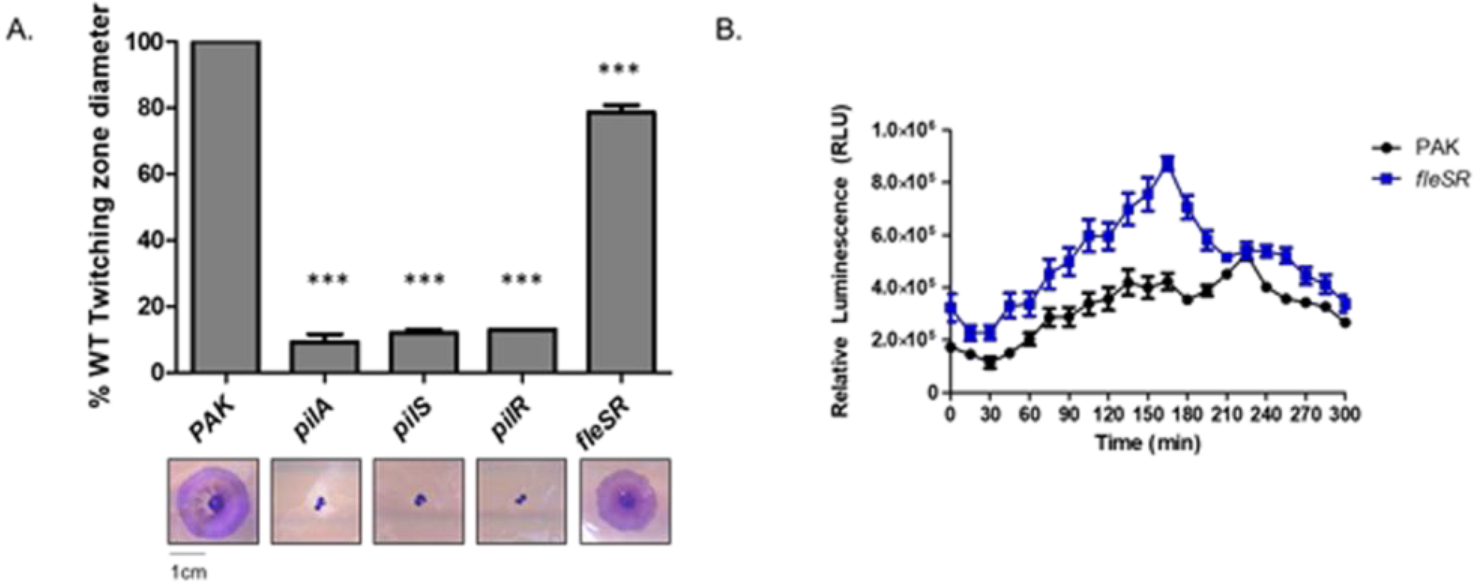
Loss of *fleSR* reduces twitching motility but increases *pilA* transcription. A) Loss of *fleSR* reduces twitching motility by approximately 20%. Mean and standard error of 6 independent replicates are shown. Significance determined by one-way ANOVA. **B)** A *lux-pilA* luminescent reporter assay measuring *pilA* promoter activity indicated that *pilA* transcription is increased over 5h. Mean and standard error of 4 biological replicates are shown.

## DISCUSSION

Two-component systems control a multitude of phenotypes, allowing for quick responses to sudden changes in a bacterium’s intra-and extracellular environments. These systems can be important for survival, but also for coordinating virulence programs. Most TCSs explored to date control the transcription of multiple genes, but prior to this work *P. aeruginosa* PilR had only a single known target, *pilA* (23). Microarray and bioinformatics analyses of the *G. sulfurreducens* PilR regulon provided evidence that PilR regulates multiple genes, including those required for soluble Fe(III) uptake (a pilin-independent phenotype), flagellar assembly and function, and cell envelope biogenesis, though these predictions were not confirmed with phenotypic assays (16,25). Here, we showed that PilR controls the expression of multiple genes, in pilin-responsive or unresponsive modes. Dysregulating expression of select members of the *P. aeruginosa* PilR regulon resulted in changes in swimming, swarming, and/or twitching motility, all phenotypes associated with virulence in specific hosts (27, 38–40).

The *G. sulfurreducens* and *D. nodosus* studies cited above failed to account for the confounding variable that PilA is not expressed when *pilR* is deleted. This was an important consideration in designing our RNAseq experiment (**Figure 1**), as loss of PilA results in decreased cAMP levels and by extension, downregulation of cAMP-dependent genes in the Vfr regulon, which includes a number of T4P-associated genes (28). This design also enabled us to further classify genes in the PilR regulon based on their responsiveness to pilin levels. As predicted, many of the genes that were similarly dysregulated by loss of both *pilA* and *pilR* are Vfr-dependent (28) **(Table S1).** Thus, we focused instead on those genes that were dysregulated in a PilR-dependent manner and further categorized them as pilin responsive or unresponsive.

We identified ten pilin-responsive genes with increased transcription in a *pilA* mutant but significantly decreased transcription in the absence of *pilR*, even though *pilR* mutants also lack PilA (**Figure 2**). While this expression pattern initially seemed counterintuitive, we propose that these gene products are regulated by PilS phosphorylation or dephosphorylation of PilR in response to fluctuating PilA levels. At high concentrations, PilA represses its own transcription by interacting directly with PilS in the inner membrane, promoting its phosphatase activity on PilR (22). Conversely, when PilA is absent, PilS phosphorylates PilR and *pilA* promoter activity is significantly increased, presumably in an attempt to replenish intracellular PilA pools (41) but simultaneously increasing expression of other pilin responsive genes (**Figure 2**, **Table S2**). This signalling pathway may be one way in which adherence of a pilus to a surface is detected, through transient depletion of pilin pools in the inner membrane when attached pilus filaments fail to retract.

Many genes in this pilin-responsive category encoded hypothetical proteins or were unannotated in the PAO1 and PAK genomes; the latter may encode regulatory RNAs. We used available mutants from the PA14 Tn library to determine if the pilin-responsive genes were required for normal T4P function. While all mutants tested had WT twitching motility, some had decreased swarming, and one (PA14_69560) had decreased swimming motility. The only genes in this group that were characterized previously are *hcpA* and *hcpB*, which encode proteins associated with the Type VI secretion system. They are paralogs, possibly resulting from a gene duplication event (35). This finding may represent a new link between T4P, flagellar function, and Type VI secretion, as the *hcpB* mutant had defects in both swimming and swarming. This connection further explains the swimming defects *of pi IS* and *pilR* mutants (**Figure 4**).

We also identified genes that were affected only by loss of *pilR*, independent of PilA. These genes might be modulated in response to cues detected by a different, pilin-insensitive sensor kinase that can activate PilR. Alternatively, they may already be expressed in the WT at levels such that further activation upon loss of *pilA* did not meet our 2-fold cutoff. A third possibility is that they are indirectly upregulated as a result of PilR activity on adjacent promoters. For example, among these genes were those encoding the T4P assembly ATPase PilB and the prepilin peptidase, PilD, which are contiguous with *pilC* encoding the platform protein; however, there were insufficient reads in our RNAseq analysis to accurately determine *pilC* expression levels (**Figure 2**). Based on this and previous studies, *pilBCD* are not co-transcribed (35,36). *pilB* was reported to be σ^70^ dependent (36), but our data suggest that PilR remodeling of the *pilA* promoter for transcription by the σ^54^ holoenzyme also facilitates transcription from the divergent *pilB* promoter.

Of the pilin unresponsive genes identified, the most abundant class were involved in biosynthesis, function, and regulation of the flagellum, including *fleSR* (**Figure 2**, **Table S3**) Most of the others are members of the FleSR regulon (6) suggesting they are indirectly regulated by PilR. Swimming motility of p/75 and *pilR* mutants was ~40% of WT, supporting the expression data (**Figure 4**). By carefully analyzing the swimming data, we hypothesized that suppressor mutations could overcome the defects imposed by *pilS or pilR* deletion, allowing the mutants to swim normally. Preliminary sequence analyses of these suppressors showed no mutations in *fleSR*, but it may be that mutations in *fleQ*, the promoter regions of *fleSR*, or as yet unidentified genes could increase activity or expression of *fleS-fleR*. The as-yet unidentified suppressors appear specific for flagellar function, as swarming motility (42) of the *pilS* and *pilR* mutants and the highly motile suppressors, all of which lack PilA, was comparable to that of a *pilA* mutant (**Figure S2**),

Although *pilSR* were not considered members of the FleSR regulon (6), twitching motility was modestly but reproducibly reduced to ~80% of WT in the absence of *fleSR* (**Figure 5A**), while *pilA* promoter activity was increased compared to WT (**Figure 5B**). This phenotype is reminiscent of mutations that inhibit pilus retraction, impairing twitching but increasing *pilA* transcription due to depletion of PilA subunits from inner membrane pools (11,41). During prior characterization of the FleSR regulon, two new genes (PA3713 and PA1096 *fleP*) with motility phenotypes were identified. Mutants were significantly impaired in swimming, and in the case of *fleP*, twitching motility (6). FleP was proposed to control pilus length, as when it was deleted, surface pili were significantly longer than those of WT, resulting in a form of hyperpiliation (6). Decreased *fleP* expression in our *fleSR* mutants could impair twitching motility and alter p/M expression. Because of their FleSR dependence, expression of *fleP* and PA3713 may be decreased in *pilSR* mutants; however, the reads for them in our RNAseq experiment were too low to assess this idea.

Both the PilSR and FleSR TCSs are required for full virulence of *P. aeruginosa*-reviewed in (43) - as each is involved in multiple virulence-associated phenotypes. PilSR and FleSR each contribute to surface attachment and biofilm formation (27,44), and are important for twitching and swimming motilities. Both PilSR and FleSR are required for swarming motility due to their involvement in pilus and flagellum function respectively (37, 42, 44), (**Figure S3**). Given the overlap in phenotypes controlled by PilSR and FleSR, it is perhaps not surprising that expression of the two systems may be linked. From our RNAseq analysis and subsequent phenotypic assays, we propose a model in which PilSR positively regulates *fleSR* transcription, independently of PilA depletion (**Figure 6**). The hierarchy for flagellar biosynthesis proposed by Dasgupta *et al.* (6) suggests that transcription of *fleSR* is predominantly dependent on FleQ. Since *fleQ* was not differentially expressed in *pilR*, we infer that PilSR promotes *fleSR* transcription directly, rather than by modulating FleQ expression.

**Figure 6:**
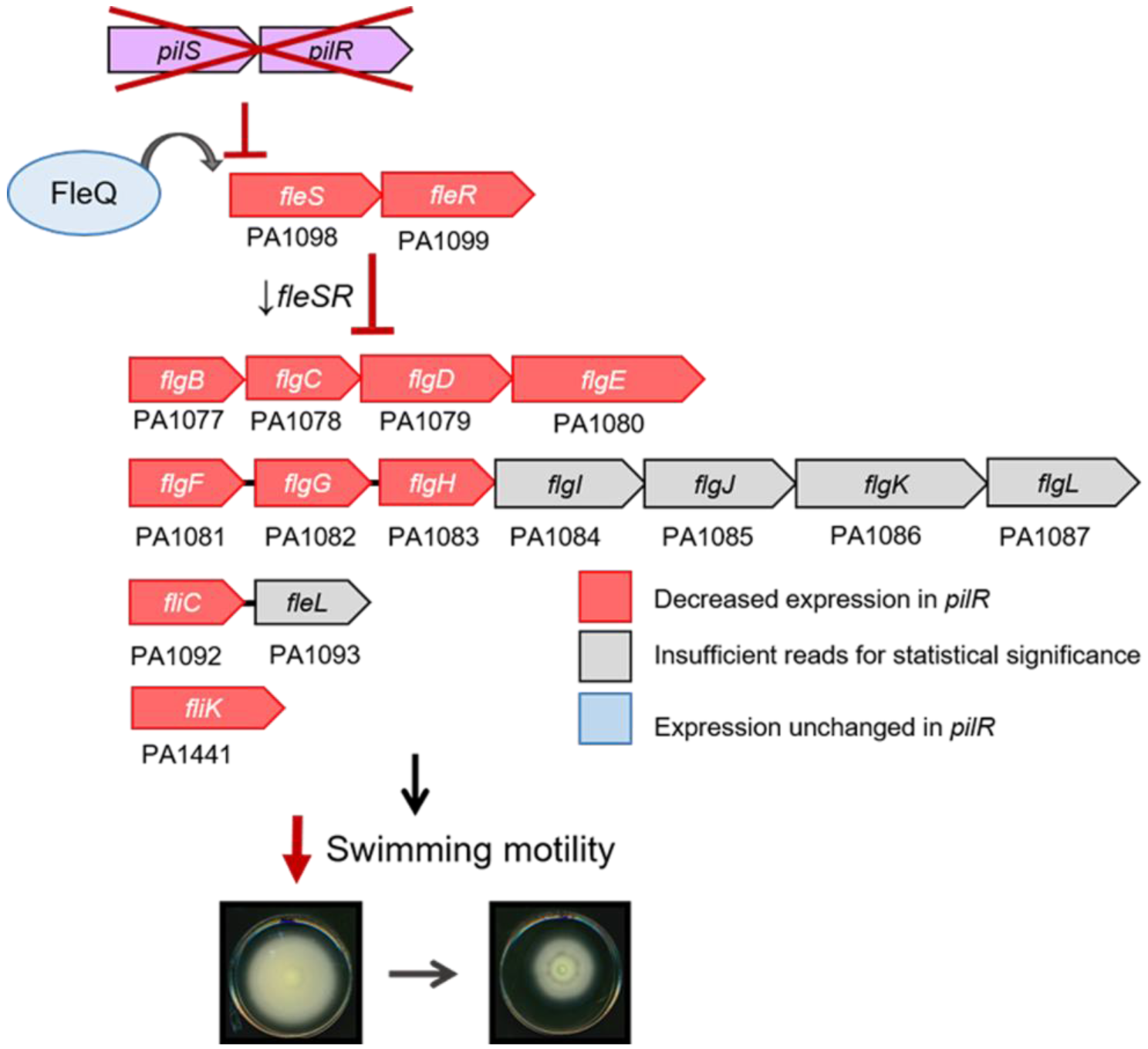
Model for *pilSR* dependent regulation of *fleSR* and the *fleSR* regulon. Under conditions in which *pilSR* expression is decreased (low cAMP) or perhaps when their activity is low (high intracellular PilA), *fleSR* transcription is decreased. As a result, in *pilSR* mutants, as is expression of the the *fleSR* regulon. Genes in red are those that had decreased expression in a *pilR* mutant in RNAseq. Grey genes did not have sufficient reads assigned to them from RNAseq to accurately report differential expression. FleQ (blue) was not differentially expressed between WT or *pilR* indicating that *pilSR* fits into the flagellar regulatory hierarchy after FleQ but before *fleSR*, as *fleQ* itself, and most FleQ dependent genes were unaffected by loss of *pilR*.

Why, and under what conditions, might this regulatory circuit be active? Twitching motility is normally deployed on solid or semi-solid surfaces (8) while flagella are typically used in liquid and low viscosity conditions. One might predict that the systems are differentially activated in response to relevant environmental conditions. Instead, the regulatory integration of these two systems may be an adaptation to life as an opportunistic pathogen. T4P and flagella are typically expressed during the acute phase of infection (4,45) and during the transition to the chronic infection phase, motility systems are downregulated in favour of those promoting Type VI secretion and biofilm formation (43,46). Clinical isolates of *P. aeruginosa* from chronically colonized patients are often both non-flagellated and non-piliated (47). Lack of the immunogenic flagellum may help *P. aeruginosa* escape phagocytosis (47) and aflagellate bacteria are better able to evade the inflammatory response of the host (48). Placing *fleSR* under control of PilSR may facilitate a more rapid transition to the chronic disease state and more efficient evasion of the host immune system. Similarly, both T4P and flagella are required for surface sensing and surface-associated behaviours such as swarming motility and activation of virulence cascades (26, 42, 49, 50). Co-regulation of their expression may allow *P. aeruginosa* and other motile bacteria to amplify their responses to surface detection.

We identified 34 genes in addition to *pilA* whose expression was altered ≥3 fold by loss of *pilR*, 24 of which were dysregulated in a pilin unresponsive manner, supporting previous work in *G. sulfurreducens* that identified putative PilR binding sites upstream of multiple genes (25). Importantly, while *pilA* and *pilR* mutants look similar with respect to their T4P-related phenotypes, their transcription profiles and other phenotypic outputs are different. For example, expression of genes encoding proteins involved in flagellum biosynthesis, including *fleSR*, are downregulated in the absence of *pilR* but unaffected by loss of *pilA*. This work reveals a previously unappreciated regulatory connection between two diverse motility systems, with implications in detection of surface attachment and the transition from acute to chronic disease states in a host.

## METHODS

### Bacterial strains and growth conditions

Unless otherwise specified, *Pseudomonas aeruginosa* PAK strains were grown in Lennox Broth (LB) (Bioshop) or on LB 1.5% agar plates at 37°C. Where the antibiotic kanamycin was used, it was introduced at a final concentration of 150ng/mL. Mutants were generated by homologous recombination, using standard mating techniques described in (51). The strains and plasmids used in this study are outlined in **Table 1.** Plasmids were prepared using standard cloning techniques and introduced into *P. aeruginosa* using electroporation.

### RNA isolation, library preparation, cDNA sequencing and analysis

To isolate RNA, cells from strains of interest were streaked in triplicate onto half of an LB 1.5% agar plate (100x15mm petri dishes) and grown overnight at 37°C. Cells were scraped from the plates and resuspended in 1.5mL RNAprotect Bacteria Reagent (Qiagen) to maintain integrity of isolated RNA. Cells were chemically lysed using 1mg/mL lysozyme in lOmM Tris-HCI and ImM EDTA, pH 8.0 and RNA isolated using the RNeasy mini kit (Qiagen) according to manufacturers’ instructions. An on-column DNase treatment was performed to minimize potential DNA contamination. Purified RNA was eluted into 50nL nuclease free water and quantified.

The following steps were performed by the Farncombe Metagenomics Facility (McMaster University, Hamilton, ON, Canada). For RNAseq analysis, ribosomal RNA was depleted from 9 RNA samples (3x WT PAK, 3x *pilA* and 3x *pilR*) using the Ribo-zero rRNA depletion kit (Illumina) and cDNA libraries prepared by the NEBnext Ultra Directional Library Kit. Libraries were sequenced using paired end 75bp reads on the Illumina MiSeq platform. Reads were aligned to the PAO1 reference genome with 98% of reads mapped and normalization and differential gene expression were calculated using the Rockhopper software (52). q-values for each identified gene are reported in **Tables S1-3.** The complete RNAseq dataset has been deposited in NCBI GEO (Accession number: GSE112597).

### Twitching motility assays

Twitching motility assays were performed as described in (53). Briefly, strains of interest were stab inoculated to the bottom of an LB 1% agar plate with a P10 pipette tip and plates were incubated upside down at 37°C for 16–24h. Following incubation, agar was carefully removed and the plastic petri dish was stained with 1% crystal violet for 20min. Excess dye was washed away with water and twitching zone diameters were quantified using ImageJ ((http://imagei.nih.gov/ii/. NIH, Bethesda, MD). A one-way ANOVA statistical test was used to determine significant differences in twitching compared to WT.

### Swarming motility assays

Swarming motility assays were performed as described in (54). Briefly, strains of interest were grown overnight in 5mL LB cultures at 37°C. On the day of the assay, 0.5% agar plates with M8 buffer,supplemented with 2mM MgS0_4_, 0.2% glucose, 0.05% L-glutamic acid and trace metals, were prepared and allowed to solidify at room temperature for 1.5h. Then, 3.5nL of culture were spotted onto the centre of a single plate and plates were incubated upright in a humidity-controlled 30°C incubator for 48h. Plates were imaged using a standard computer scanner. Figures shown are representative of 3 independent experiments.

### Swimming motility assays

Swimming motility plate assays were performed similarly to (55), with some modifications. Overnight 5mL cultures of strains of interest were grown at 37°C in LB with shaking. On the day of inoculation, LB0. 25% agar plates were prepared and allowed to solidify at room temperature for 1.5h. Cell cultures were standardized to an OD_600_=1.0 and 2μL were spotted onto the centre of each plate. Plates were incubated upright for 16h at 37°C and swimming zone diameters were quantified using ImageJ (http://imagei.nih.gov/ii/. NIH, Bethesda, MD). Where applicable, swimming zone diameters were defined at the outer most part of the swimming zone that was still uniform in appearance. Images are representative of 4 independent experiments. To determine statistical significance, a one-way ANOVA analysis with Dunnett’s post-test was performed, using WT as the control strain.

### Biofilm assays

Biofilm assays were performed similarly to the method described in (33), with some modifications. Briefly, *P. aeruginosa* strains of interest were grown in 5mL liquid cultures of 50% LB/50% PBS (50/50 media) overnight at 37°C with shaking. The following day, strains were subcultured 1:25 into fresh 50/50 media and grown to a standardized OD_600_=0.1. Standardized cultures were then diluted 1:500 and 150nL of each strain of interest was plated in triplicate in a clear, 96 well plate (Nunc). The plate was closed with a 96-peg lid, providing a surface on which biofilms can form, sealed with parafilm and incubated with shaking for 18h at 37°C. To quantify planktonic growth, peg lids were removed and the 96-well plate was scanned at a wavelength of 600nm. To quantify biofilms, peglids were washed in PBS and stained with crystal violet for 15min. Following five lOmin washes in water, crystal violet was solubilized in 33% acetic acid in a fresh 96 well plate, which was scanned at 595nm. Biofilm data was graphed as % WT, showing means and standard error of three independent experiments.

### Caenorhabditis elegans slow killing pathogenicity assays

Slow killing (SK) assays were performed as described previously (34). *Caenorhabditis elegans* strain N2 populations were propagated and maintained on Nematode Growth Media (NGM) plates inoculated with *E. coli* OP50. Eggs were harvested to obtain a synchronized population by washing worms and eggs from NGM plates with M9 buffer. Worms were degraded by adding buffered bleach, leaving only eggs intact. Eggs were washed with M9 buffer and resuspended in M9 buffer with rocking overnight to allow eggs to hatch into L1 larvae. Synchronized L1 worms were plated on NGM plates for 45h to develop into L4 worms. During this process, slow killing plates supplemented with lOOμM 5-Fluoro-213-deoxyuridine (FUDR) were prepared and inoculated with lOOμL of a 5mL LB overnight culture of bacterial strains of interest and incubated at 37°C for 16–18h. Harvested and washed L4 worms (~30–40) were dropped by Pasteur pipette onto each SK plate. Using a dissecting microscope, plates were scored daily for dead worms, which were picked and removed. Survival curves were prepared using Graphpad Prism 5.01 (La Jolla, CA) and statistically significant differences in pathogenicity between strains were identified using Gehan-Breslow-Wilcoxon analysis.

### pilA-lux reporter assay

Luminescent reporter assays were performed as described previously (22). Strains of interest were transformed by electroporation with the pMS402-p*pilA* plasmid, which contains the luciferase genes under control of the *pilA* promoter. Strains were grown overnight in 5mL LB cultures supplemented with 150 ng/mL kanamycin. The following day, a 1mL aliquot of a 1:20 dilution of cultures was prepared and lOOμL samples were plated in triplicate in a white walled, clear bottom 96-well plate (3632 Costar, Corning Inc). Luminescence and OD_60_0 were measured at 15min intervals over 5h using a Synergy 4 microtitre plate reader (BioTek) programmed to shake continuously and incubate the plate at 37°C. Luminescence was normalized to OD_60_0 and relative luminescence was plotted against time. Mean and standard error of >4 biological replicates are shown.

## AKNOWLEGDEMENTS

The authors thank Stephanie Jones for assistance with the RNAseq data analysis and Michael Surette for the PA14 transposon mutant library. This work was funded by a Canadian Institutes of Health Research grant MOP 86639 to LLB. SLNK held an Ontario Graduate Scholarship.

**Figure SI: RT-PCR validation of *fleSR* transcription levels**. *fleS-fleR* were downregulated approximately 2-fold in *pilS* and *pilR*, but not *pilA*. Mean and standard error of 3 independent experiments are shown.

**Figure S2: Putative suppressors that restore swimming do not affect swarming motility in** *pilSR* mutants. Cells from *pits* and *pilR* mutants with putative suppressor mutations that restore swimming motility (outer) were inoculated in a swarming assay for comparison to *pilS* and *pilR* mutants that do not have suppressors (inner). Strains with possible suppressors still exhibit *pilS* and *pilR*-like swarming motility patterns indicating that the *pilSR* phenotype is dominant.

**Table SI. Genes similarly dysregulated in *pilA* and *pilR***

**Table S2. Genes inversely dysregulated in *pilA* and *pilR***

**Table S3. Genes dysregulated by loss of *pilR* only**

